# Developmental encoding of ultrasound vocalizations in the mouse auditory cortex

**DOI:** 10.1101/2024.07.05.601869

**Authors:** Stefano Zucca, Chiara La Rosa, Tommaso Fellin, Paolo Peretto, Serena Bovetti

**Affiliations:** Department of Life Sciences and Systems Biology (DBIOS), University of Turin, via Accademia Albertina 13, 10123, Torino, Italy; Neuroscience Institute Cavalieri Ottolenghi (NICO), University of Turin, Regione Gonzole 10, 10143, Orbassano, Italy; Optical Approaches to Brain Function Laboratory, Istituto Italiano di Tecnologia, via Morego, 30, 16163 Genoa, Italy

**Author notes:** Corresponding author: Serena Bovetti, PhD, University of Turin, Dept. of Life Sciences and Systems Biology (DBIOS), Via Accademia Albertina, 13-10123 Torino-IT, Neuroscience Institute Cavalieri Ottolenghi (NICO), Regione Gonzole, 10-10043 Orbassano (To) – IT.

**Keywords:** auditory cortex, mouse communication, spontaneous activity, two-photon calcium imaging, ultrasound vocalizations

## Abstract

Mice communicate through high-frequency ultrasound vocalizations (USVs), which are crucial for social interactions such as courtship and aggression. Although USV representation has been found in adult brain areas along the auditory pathway, including the auditory cortex (ACx), no evidence is available on the neuronal representation of USVs early in life. Using in vivo two-photon calcium imaging, we analyzed ACx layer 2/3 neuronal responses to USVs, pure tones (4-90 kHz), and high-frequency modulated sweeps from postnatal day 12 (P12) to P21. We found that ACx neurons are tuned to respond to USV syllables as early as P12-P13, with an increasing number of responsive cells as the mouse age. By P14, while pure tone responses showed a frequency preference, no syllable preference was observed. Additionally, at P14, USVs, pure tones, and modulated sweeps activate clusters of largely non-overlapping responsive neurons. Finally, we show that while cell correlation decreases with increasing processing of peripheral auditory stimuli, neurons responding to the same stimulus maintain highly correlated spontaneous activity after circuits have attained mature organization, forming neuronal sub-networks sharing similar functional properties.

## Introduction

Mice communicate through a variety of signals that include the emission of complex ultrasound vocalizations (USVs), spanning a range of 35 to 110 kHz (Holy and Guo, 2005). USVs display complex acoustic features, suggesting that multiple categories of vocalizations exist and differently influence distinct social behaviors (Sangiamo et al., 2020). Mouse vocalizations consist of multiple syllables, which are continuous sound units separated by brief periods of silence, organized into nonrandom patterns, often repeated in phrases (Agarwalla et al., 2023; Arriaga and Jarvis, 2013; Holy and Guo, 2005). Changes in the sequence of syllables, their acoustic properties, and prevalence within a phrase provide insights into specific emitter characteristics such as developmental stage, genetic strain, health, and social status. Therefore, USVs play a fundamental role in many mouse social behaviors such as courtship, mate selection, and aggression (Chabout et al., 2015; Fonseca et al., 2021; Grimsley et al., 2011; Lenschow et al., 2022; Scattoni et al., 2011). Responses to USVs have been characterized in different adult rodent brain areas, including the primary auditory cortex (ACx), as well as in higher-order ACx regions (Calhoun et al., 2023; Carruthers et al., 2013; Geissler and Ehret, 2004; Levy et al., 2019), and both mice and rats have been shown to overexpress USV frequency in the auditory pathway (Afrashteh et al.,2022; Garcia-Lazaro et al., 2015; Kim and Bao, 2009), supporting the importance of encoding ethologically relevant sounds.

In altricial species, the capacity to respond to auditory stimulation develops postnatally with the opening of the auditory canal occurring around P11 in mice (Chen et al., 2021; Polley et al., 2013), although wide-field calcium imaging revealed responses before ear canal opening (Babola et al., 2018; Meng et al., 2020). Pure tone tonotopic responses have been shown to develop rapidly between P12 and P14, reaching an adult-like representation at P14-P16 in mice (Carrasco et al., 2013; Chen et al., 2021; Polley et al., 2013). The early development of tonotopy comes with a marked increase in dendritic complexity and spine number, which is followed by a period of spine maturation approximately after P16 (McMullen et al., 1988; Meng et al., 2020; Schachtele et al., 2011), which corresponds to the functionally identified critical period for audition (Barkat et al., 2011; Meng et al., 2020; Zhang et al., 2001).

The development of complex sound responses is slightly delayed compared with pure-tone responses and emerges in a series of sensitive periods within a month-long critical period window (Carrasco et al., 2013; Insanally et al., 2010; Insanally et al., 2009; Kim and Bao, 2013). Interestingly, early exposure to complex ultrasound vocalization frequencies is required for high-frequency (HF, > 45KHz) overrepresentation in the rat auditory cortex, suggesting that early experience has a significant impact on complex sound processing at both neuronal representational and behavioral levels.

Importantly, the maturation of auditory circuits is influenced by “spontaneous”, i.e. sensory-independent, neuronal activity. Action potentials that are not initiated by input from the external environment have been described in ACx in a period starting shortly after birth and lasting until hearing onset at the opening of the auditory canal (Wang and Bergles, 2015), when stimuli-evoked responses initiate shaping the organization of nascent circuits with consequent increase of cell desynchronization (Meng et al., 2021; Meng et al., 2020; Wang and Bergles, 2015). Using macroscopic calcium imaging on both the inferior colliculus and auditory cortex before hearing onset, it has been shown that neurons responsible for processing similar frequencies of sound exhibit highly synchronized activity at this early developmental age (Babola et al., 2018), providing a mechanism for the activity-dependent refinement and stabilization of synaptic connections within specific frequency ranges.

Despite the prominent role of USVs in mouse social behaviors and communication, no evidence is available on the neuronal representation of USVs early in life. Previous studies have mainly focused on ACx tonotopy development and neuronal representation of pure tones (< 32 kHz). However, conspecific vocalizations differ from pure sounds not only for their complex spectrotemporal structure but also for their natural incentive salience (Sangiamo et al., 2020).

How and when does the capacity to respond to a USV presentation develop? Are different syllables composing a USV bout differently represented in the developing ACx?

In the present study, we analyzed layer 2/3 (L2/3) ACx representation of USVs from hearing onset (∼P12 in mice) to P21 and compared responses to isolated syllables from conspecific USVs with pure tones (ranging from 4 to 90 kHz) and with up-(70 – 90 kHz) and down-(90 - 70 kHz) HF-modulated sweep presentation. By performing in vivo two-photon Ca2+ imaging in anesthetized mice in four developmental windows (P12-P13, P14-P15; P18-P19, P20-P21) we characterize single-cell responses to different sound categories and selective stimuli. Moreover, we examined spontaneous activity and network correlation at different ages and evaluated the relationship between spontaneous and sound-evoked activity.

Our results show that ACx L2/3 neurons are tuned to respond to USV syllables as early as P12-P13. The fraction of neurons that respond to single syllable increases during development; however, contrary to pure-tone responses, we did not detect age-related preference for a specific syllable feature. Finally, through the analysis of pairwise cell correlations of both spontaneous and evoked neuronal activity, we demonstrate that clusters of neurons involved in processing similar simple (e.g. pure tones) or complex (e.g. syllables) sounds display higher correlated spontaneous activity after the opening of the auditory canal, coinciding with the initiation of sensory-evoked network activation.

## Materials and Methods

### Animals

Experimental procedures involving animals were approved by the University of Turin ethical Committee and by the National Council on Animal Care of the Italian Ministry of Health (authorization # 422/2022-PR). All experiments were conducted according to the guidelines of the European Communities Council Directive of November 24, 1986 (86/609/EEC). The animals were housed under a 12-h light: dark cycle in individually ventilated cages. Two-photon imaging experiments were performed on C57BL/6j mice (either males or females) for 12 days after birth (P12) to 21 days (P21). Breeding cages were organized with a single adult male bred with one or two adult females. Newborn pups were kept in the cage until the day of the experiment.

### Viral Injection

To express the calcium indicator GCaMP in the neurons of the auditory cortex, the adeno-associated virus (AAVs) pGP-AAV-syn-jGCaMP7s-WPRE (AAV1; Addgene) was injected (1:10 in saline) in newborn mice at P0 – P1 as previously reported (Zucca et al., 2017). Briefly, P0-P1 mice were deeply anesthetized by hypothermia, placed on a custom-made stereotaxic apparatus, and kept at 4°C throughout the entire surgery. The skull was exposed by a small skin incision over the auditory cortex, and ∼250nl of viral suspension was injected using a glass capillary at stereotaxic coordinates of 1 mm posterior and 1.5 mm lateral to the bregma and 0.1–0.2 mm depth to target superficial layers. The capillary was kept in place for 2 min before retraction. The skin was then sutured, and the pup was revitalized under an infrared heating lamp.

### Ultrasound vocalization recording

Ultrasound vocalizations were recorded from two adult male C57BL/6j mice following acute exposure to conspecific female odors. To elicit reliable USVs, male mice were first exposed to female mice for at least 3 consecutive days. On the fourth day, male mice were exposed only to female scent marks. Scent marks consisted of a mixture of nesting and bedding materials from a single cage containing at least two adult female mice. Females were housed in clean cages, and scent marks were freshly collected the morning of the following day. To record USVs, male mice were first habituated to the soundproof recording chamber in their cage for 15 min, after which the female stimulus was placed in the cage and USVs were recorded using a calibrated ultrasound microphone (UltraSoundGate CM16/CMPA, Avisoft) for a total time of 15 min.

### Acoustic Stimuli

Sound-evoked responses were triggered using a set of custom-made stimuli. Pure Tones (Fig. 1 and Fig. 3), consisted of a sequence of five 500-ms-long sound stimuli at a single frequency ranging from 4 kHz to 64 kHz and spaced one octave each (4 kHz, 8 kHz, 16 kHz, 32 kHz, 64 kHz), with a 10-ms fade-in and fade-out window. The five different pure tones were randomly presented10n times each, with an interstimulus interval of 5 s. Syllable stimuli (Fig. 1 and Fig. 2) were generated by selecting five different syllables from the USV recordings of two male C57BL/6j adult mice. The full USV recorded signal was first filtered with a high-pass filter at 40 kHz, and noise reduction was applied to the filter trace (threshold: -70 dB, reduction: -140 dB). Syllables were cut from cleaned USV traces for a total length of 30 ms each. Each stimulus consisted of five repetitions of the same syllable spaced 70 ms each, resulting in a total stimulus length of 500 ms. All five syllables were presented10n times in a random sequence with an interstimulus interval of 5 s. High-frequency pure tones (Fig. 4) were generated by concatenating five 30-ms-long pure sounds with a 2-ms fade-in and fade-out window and spaced 70 ms each for a total stimulus length of 500 ms. Three different frequencies were chosen (70 kHz, 80 kHz, 90 kHz) to cover the frequency range of USVs. In addition, up- and down-sweeps were similarly generated, ranging from 70 kHz to 90 kHz and 90 kHz to 70 kHz,, respectively. Each sweep lasted 30 ms with a 2-ms fade-in/out and was repeated five times with 70-ms intervals, for a total stimulus length of 500 ms. Both High-Frequency and Sweep stimuli were presented 10 times in a random sequencewithh an interstimulus interval of 5 s. Audio files (.wav) were generated using a custom-made MatLab script.

### Two-photon calcium imaging

To collect sound-evoked responses in layer 2/3 neurons of the mouse auditory cortex, two-photon calcium imaging was performed on young (P12-P21) mice injected with the AAVs expressing the calcium indicator GCaMP7s at P0 – P1. Mice were first deeply anesthetized with a mixture of Ketamine/Xylazine (50mg/kg, 4mg/kg intraperitoneal administration). The skin was then removed to expose the skull, and a 3D printed head-plated was attached to the skull with surgical glue (VetBond) and dental cement, centered over the coordinates of the ACx (1 - 2mm AP from Bregma, 3 - 4mm ML from midline, depending on age). A small craniotomy (∼ 300 x 300 μm) was opened on top of the ACx, carefully maintaining the dura intact. The surface of the brain was kept moist with normal HEPES-buffered artificial cerebrospinal fluid, and body temperature was maintained at 37 °C with a heating pad. The depth of anesthesia was monitored by checking the respiration rate and reactions to pinching the tail and toe. To collect calcium signals, animals were placed under a standard laser scanning two-photon microscope (Nikon A1RMP) coupled to a Chameleon Ultra II (Coherent Santa Clara, CA, λexc = 920 nm). The activity of layer 2/3 neurons was monitored by imaging GCaMP fluorescence via a 16x objective (Nikon CFI75 LWD 16xW NIR A.N.0,80 d. l. 3,0 mm). Signals were acquired by collecting temporal series (t-series) images at an acquisition frame rate of 4 Hz and laser power ranging between 20 and 50 mW. Sound responses were evoked by placing an ultrasound speaker (Ultrasonic Speaker Vifa, Avisoft) connected to an ultrasound playback interface (UltraSoundGate Player 116H) controlled by a digital trigger (TTL) from the two-photon system. The speaker was placed approximately 10 cm from the contralateral right ear of the mouse, and neuronal activity was collected from the left ACx. Stimulus intensity was calibrated by placing the ultrasound microphone in the same position as the animal under a two-photon microscope.

### Rhodamine labeling of the recorded region for ACx anatomical confirmation

The fluorescent dye Rhodamine was used to check for correct anatomical localization of two-photon recordings in ACx. At the end of each experiment, a glass capillary was stained with rhodamine and gently inserted into the imaged craniotomy. Each mouse was then sacrificed, and the brains were extracted and left in a 4% Paraformaldehyde in PBS (pH 7.4) overnight for tissue fixation. Fixed brains were socked in a 30% sucrose PBS solution for cryoprotection, and coronal slices (50 µm-thick) were later cut and sequentially collected. Slices were stained with DAPI (1:1000, 20 min at RT), mounted on a glass slide with Mowiol mounting medium, and images were acquired via a standard confocal microscope. ACx anatomical confirmation was assessed by checking the rhodamine fluorescent signal along the capillary track in slices aligned with the reference mouse brain atlas (Kronman et al., 2023; https://kimlab.io/brain-map/epDevAtlas/) for ACx localization.

### Two-photon Analysis: Sound-evoked responses

To assess sound-evoked responses, fluorescent signals from layer 2/3 neurons imaged via two-photon microscopy were extracted using the Python-based software Suite2P (Pachitariu et al.,2017). T-series images from the same Field of View (FOV) were concatenated and Regions of Interest (ROIs) corresponding to single neurons were automatically identified and manually checked. For each identified ROI, the fluorescent signal and neuropil signal were extracted and stored for subsequent analysis using a custom-made MatLab script. Each signal was first adjusted for neuropil contamination by subtracting 70% of the neuropil signal from the raw cell fluorescence signal (Chen (Chen et al., 2013), and the dFF was calculated as follows:

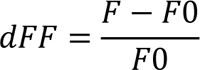

Where F is the raw fluorescent signal corrected for neuropil contamination and F0 is the median value of the raw fluorescence calculated across the entire recording.

To characterize sound-evoked responses, calcium traces were aligned to the stimulus presentation, and a time window of 5 s before and after stimulus onset was considered. Neuronal responsiveness was assessed following a previously applied method (Schiavo et al., 2020). Briefly, for each neuron, all 10 repetitions of the same stimulus were considered, and the area under the fluorescence curve was calculated for each trial considering 1 s after stimulus onset (POST) with a 250-ms blank window and 1 s before stimulus onset (PRE). A neuron was classified as responsive if there was a significant difference by comparing the PRE and POST values of the fluorescence area using the parametric paired t-student test (two tails) and if the average response calculated in the POST window was higher than the 1.5*STD calculated in the PRE window. The fraction of responsive neurons was calculated for each single animal as responsive neurons across all FOVs divided by the total number of cells recorded from the same animal. To evaluate the best frequency response, only cells responsive to at least one pure tone were considered, and the stimulus with the highest average dFF peak response was considered as the best frequency.

### Two-photon Analysis: Spontaneous Activity

To evaluate the spontaneous activity of layer 2/3 neurons and the relationship between spontaneous and sensory-evoked activity, the basal neuronal activity for each neuron was collected in a time window of 1 min before the start of acoustic stimulation. The raw fluorescence was extracted for each single cell using the method described above. To evaluate the correlation coefficient across paired cells, the Pearson correlation coefficient was calculated considering the whole basal neuronal activity across all neurons recorded within the same FOV. The distribution of all correlation coefficients was calculated for each single animal and averaged across animals of the same age group. To evaluate how the correlation coefficient changes as a function of neuronal distance, we calculated the distance between paired neurons of the same FOV by considering the centroid coordinates obtained from the ROIs identified by Suite2P. Cell distance was then binned (bin size: 10 μm), and the average correlation coefficient was calculated by averaging the correlation coefficients of cells from the same animal with a distance within the distance bin range. Distributions were calculated and averaged across FOVs from animals of the same age range. To assess the average correlation across responsive and nonresponsive cells, cells of the same FOV sharing the response for the same stimuli were pooled together and their correlation coefficient for spontaneous activity was calculated. Quantification was performed considering each cell across all FOVs and all animals.

### Statistics

The exclusion/inclusion criteria were based only on technical and anatomical issues. Specifically, recordings with technical issues (e.g. movement during calcium imaging recordings) or in which a posteriori analysis of rhodamine staining resulted to be non-centered on the auditory cortex were excluded from the analysis. All recordings with no technical issues and anatomically centered on the auditory cortex were included.

Graphs are represented using both the median and/or the average values and showing maximum, minimum, and outlier values, with the exception of Figs. 4B, 5A, and 5B, where values are expressed as average ± sem. Non-parametric Kruskal-Wallis ANOVA and Wilcoxon signed rank tests were used when comparing median values. Statistical analysis was performed using MatLab or OriginLab software.

## Results

### Layer 2/3 neurons in the auditory cortical area respond to high-frequency complex syllables at hearing onset

To investigate the response of L2/3 neurons to USVs during development, we performed in vivo two-photon functional imaging in the auditory cortex of P12–21 C57BL/6j mice. We exposed them to a subset of syllables isolated from male mouse USVs and pure tones, and quantified the number of responsive cells as well as the proportion of neurons selectively responding to syllables, pure tones, and both stimuli (Fig. 1). We focused on four developmental windows (Fig. 1A-C), starting from the time window that follows the opening of the auditory canal (Anthwal and Thompson, 2016; Ehret 1983; Mikaelian and Ruben, 1965) up to P21, corresponding to the functionally identified late critical period for audition (Jouhaneau and Bagady, 1984; Meng et al., 2020; Yang et al., 2012; Zhang et al., 2001).

P0 mice were injected in the left ACx (Calhoun et al., 2023) with AAVs driving the GCaMP7s calcium indicator under the hSynapsin promoter (Dana et al., 2019; Fig. 1D), and both their spontaneous activity and sound-evoked responses were recorded. Sound stimuli consisted of the playback of syllables isolated from conspecific USVs (syllable 1: short; syllable 2: step down; syllable 3: up-frequency modulation; syllable 4: flat; syllable 5: step down. Frequency range: 75–100 kHz; Fonseca et al., 2021). For syllable presentation, we used a sound level between 55 and 65 dB, which we show was the intensity of male emitted USVs (Supplementary Fig. 1). On the same field of view, the response to pure tones presentation was also recorded (4 kHz, 8kHz, 16kHz, 32kHz and 64kHz, sound intensity = 70-80 dB; Romero et al., 2020). At the end of the recording phase, rhodamine was injected into the central portion of the craniotomy and all brains were withdrawn. For each brain, we confirmed that the recordings were performed in the target area by analyzing *a posteriori* rhodamine expression (Fig. 1E, see Material and methods for details).

We first defined responsive cells as any cell showing both 1) an average response to 10 presentations of the same stimulus above a threshold set at 1.5 standard deviations of the baseline averaged activity and 2) a significant increase in the area under the fluorescent curve after stimulus onset (Schiavo et al., 2020; see Materials and methods). On the total number of detected cells (Fig. 1D, bottom panel), we found that the percentage of cells responding to an auditory stimulus (either syllables and/or tones) increased during development (Kruskal-Wallis ANOVA, p = 0.00776) from 13% at P12-P13 (median; N = 5 animals) to 17% at P14-P15 (median; N = 13 animals), 37% at P18-P19 (median; N = 6 animals), and 51% at P20-P21 (median; N = 6 animals; Fig. 1B), consistent with previous calcium imaging and microelectrode recordings (Meng et al., 2020; Zhang et al., 2001). We next evaluated the proportion of responsive cells selectively activated by syllables or pure tones and the proportion of cells responding to both types of stimuli in different developmental windows. The percentage of cells responding only to syllables or tones increased during development from 6% and 8% at P12-P13 (median, N = 5 animals) to 15% and 21% at P20-P21 (median, N = 6 animals) for syllables and tones, respectively (Kruskal-Wallis ANOVA, p = 0.04641 for tones; p = 0.04982 for syllables; Fig. 1C). Similarly, later developmental ages (P18-P19 and P20-P21) also exhibited an increasing fraction of cells responding to both types of stimuli that set at 14% at P20-P21 (Kruskal-Wallis ANOVA, p = 0.00203). Interestingly, while responses to tones constantly increased from P12 to P21, syllable-evoked responses largely raised at P18-P21, consistent with previous data in rats, showing that ultra-sound frequency representation is slightly delayed compared with pure-tone low-frequency responses (Kim and Bao, 2013).

**Fig. 1.**
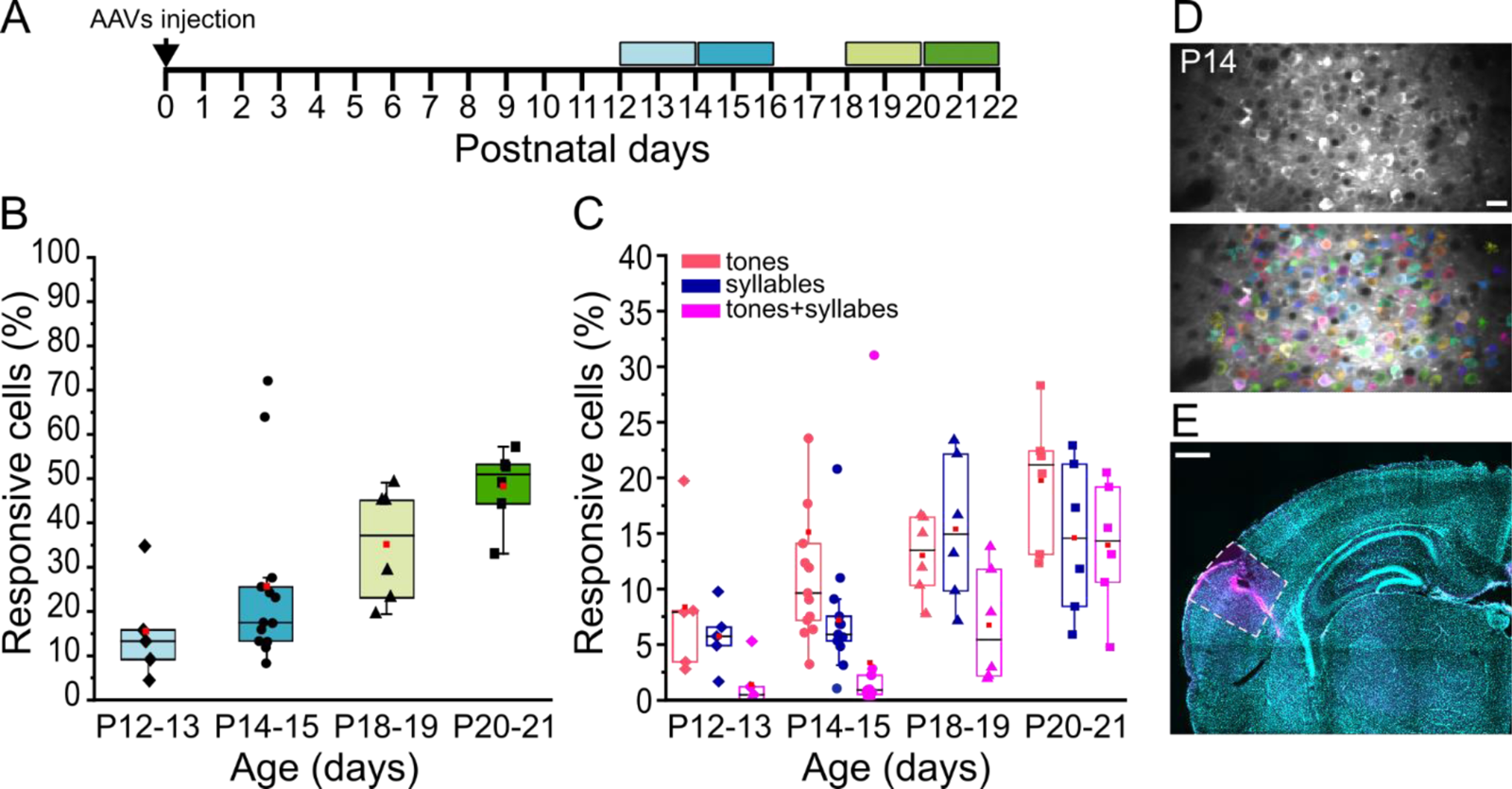
Response properties of L2/3 neurons in auditory cortical area (ACx) recorded by two-photon Ca2+ imaging. (A) Experimental protocol. P0 mice were injected in the left ACx with AAV1.hsynGCaMP7s and both spontaneous and evoked activity was recorded in four developmental windows: P12-P13, P14-P15, P18-P19 and P20-P21. (B-C) Overall fraction of responsive cells (B) and percentage of cells responding to tones (red), syllables (blue) or both stimuli (magenta) (C), in the four developmental windows. For each age the box plots show the median (horizontal black line) and the average value (red square). The overall fraction of responsive cells increases during development (B; Kruskal-Wallis ANOVA, p = 0.00776) as well as the percentage of cells responding to pure tones, syllables, or both (C; Kruskal-Wallis ANOVA, p = 0.04641 for tones; p = 0.04982 for syllables; p = 0.00203 for tones and syllables). P12-P13 N = 5; P14-P15 N = 13; P18-P19 N = 6; P20-P21 N = 6. (D) Example of a field-of-view (FOV) showing GCaMP7s expression in ACx L2/3 neurons (top panel) and same FOV with cells detected by Suite2p (Pachitariu et al.,2017) labeled with different colors (bottom panel). Scale bar = 20 µm. (E) *A posteriori* confirmation of the imaged area. Example of a coronal section from a P14 mouse brain, showing rhodamine labelling (magenta) in auditory cortical area and counterstained with DAPI (blue). Scale bar = 500 µm.

### A subpopulation of L2/3 cells in the auditory area selectively responds to complex sound features

Having observed syllable-evoked activation in the auditory cortex as early as P12-P13, we next characterized the response of single neurons in ACx L2/3 to the playback of different types of syllables during development (Fig. 2). Single syllables were isolated from USVs emitted by adult C57BL6/j male mice when exposed to female urine, and played-back at 55-65 dB using a speaker positioned 10 cm from the contralateral mouse ear (see Materials and methods for details in recording and playback procedures). Syllables were chosen on the basis of their differences in average frequency, shape, and length (Supplementary Fig. 1). To assess whether L2/3 cells in ACx are tuned to syllable identity, we first quantified the percentage of neurons responsive to one, two, or more syllables. At all developmental stages, we found that most of the neurons were selective for one syllable only (Fig. 2A-E). In particular, at P12-P13 and P14-P-15, almost all L2/3 cells responded to a single syllable, whereas at later developmental stages, the fraction of cells responsive to two or more stimuli increased (Fig. 2E). We then asked whether L2/3 cells in ACx preferentially respond to a specific syllable type at different developmental stages. We found that the number of responsive cells for each syllable type increased from P12 to P21, but we did not observe a clear preference for any of the syllable types at all ages (P12-P13: Kruskal-Wallis ANOVA, p = 0.5459, N= 5; P14-P15: Kruskal-Wallis ANOVA p = 0.152, N=13; P18-P19: Kruskal-Wallis ANOVA p = 0.1105, N=6; P20-P21: Kruskal-Wallis ANOVA p = 0.09984, N=6; Fig. 2F).

**Fig. 2.**
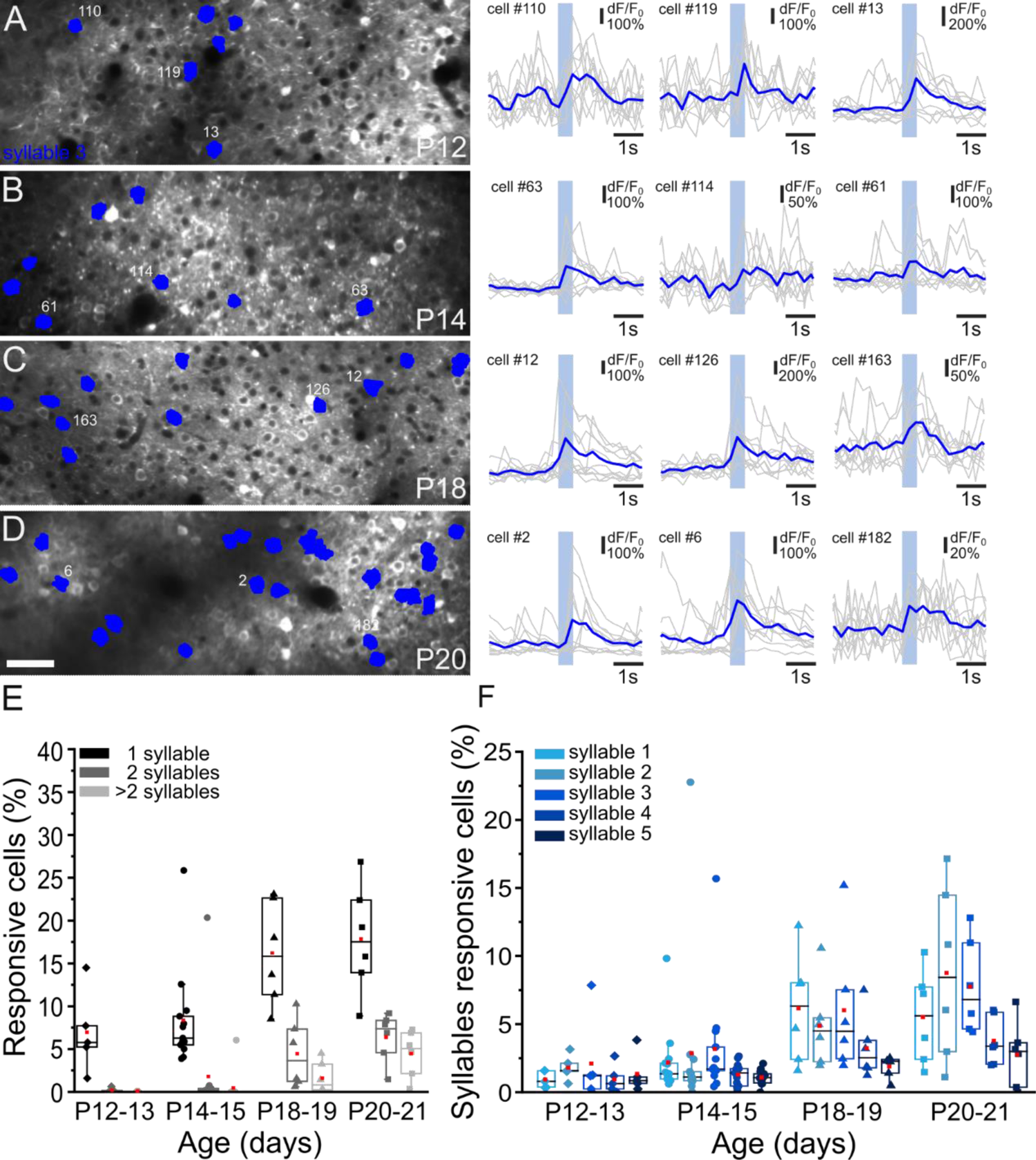
Selective responses to syllables during development. (A-D) *Left panels*: Examples of fields-of-view showing GCaMP7s expression in ACx L2/3 neurons at P12 (A), P14 (B), P18 (C) and P20 (D). Cells responding to syllable 3 are color-coded in blue. Scale bar = 50 µm. *Right panels*: Fluorescence signals (dF/F0) over time from three representative cells responding to playback of syllable 3 at different developmental ages. Gray lines represent single trial response. Blue lines represent the averaged response to ten stimulus presentation (blue rectangle). (E-F) Percentage of cells responding to one syllable, two syllables or to more then two syllables (E) and fraction of cells selectively responding to syllable 1-5 (F) in the four developmental windows. For each age the box plots show the median (horizontal black line) and the average value (red square). No preference toward a syllable was identified (Kruskal-Wallis ANOVA; P12-P13: p = 0.5459, N= 5; P14-P15: p = 0.152, N=13: P18-P19: p = 0.1105, N=6; P20-P21: p = 0.09984, N=6).

To investigate if L2/3 selectivity was restricted to syllables or was a common feature of sound responses, we characterized the responses of all recorded cells to each of the five pure tonal stimuli, ranging from 4 kHz to 64 kHz and separated by one octave each (Fig. 3). Consistent with what was found for syllable presentation, L2/3 cells were mostly selective for one stimulus only, with an increased number of cells responsive to two or more stimuli at later developmental stages (Fig. 3A-E). When looking at the preference for a single tone, at P12-P13 we did not detect a prevalent response to a specific tone (Kruskal-Wallis ANOVA, p = 0.96504, N= 5; Fig. 3F). In contrast, in P14-P15 old mice, L2/3 cells showed preferred responses to low-frequency tonal stimuli between 4 and 16 kHz (Kruskal-Wallis ANOVA p = 0.02941, N=13; Fig. 3F), consistent with previous studies showing rapid changes in simple tone responses between hearing onset and P14-P16 (Carrasco et al., 2013; Chen et al., 2021; de Villers-Sidani et al., 2007; Meng et al., 2020; Polley et al., 2013). At P18-P19, recorded cells showed a shift in tonal preference toward higher frequencies, between 16 and 32 kHz (Kruskal-Wallis ANOVA p = 0.00193, N=6; Fig. 3F), whereas no differences were detected at P20-P21 (Kruskal-Wallis ANOVA p = 0.28784, N=6; Fig. 3F).

**Fig. 3.**
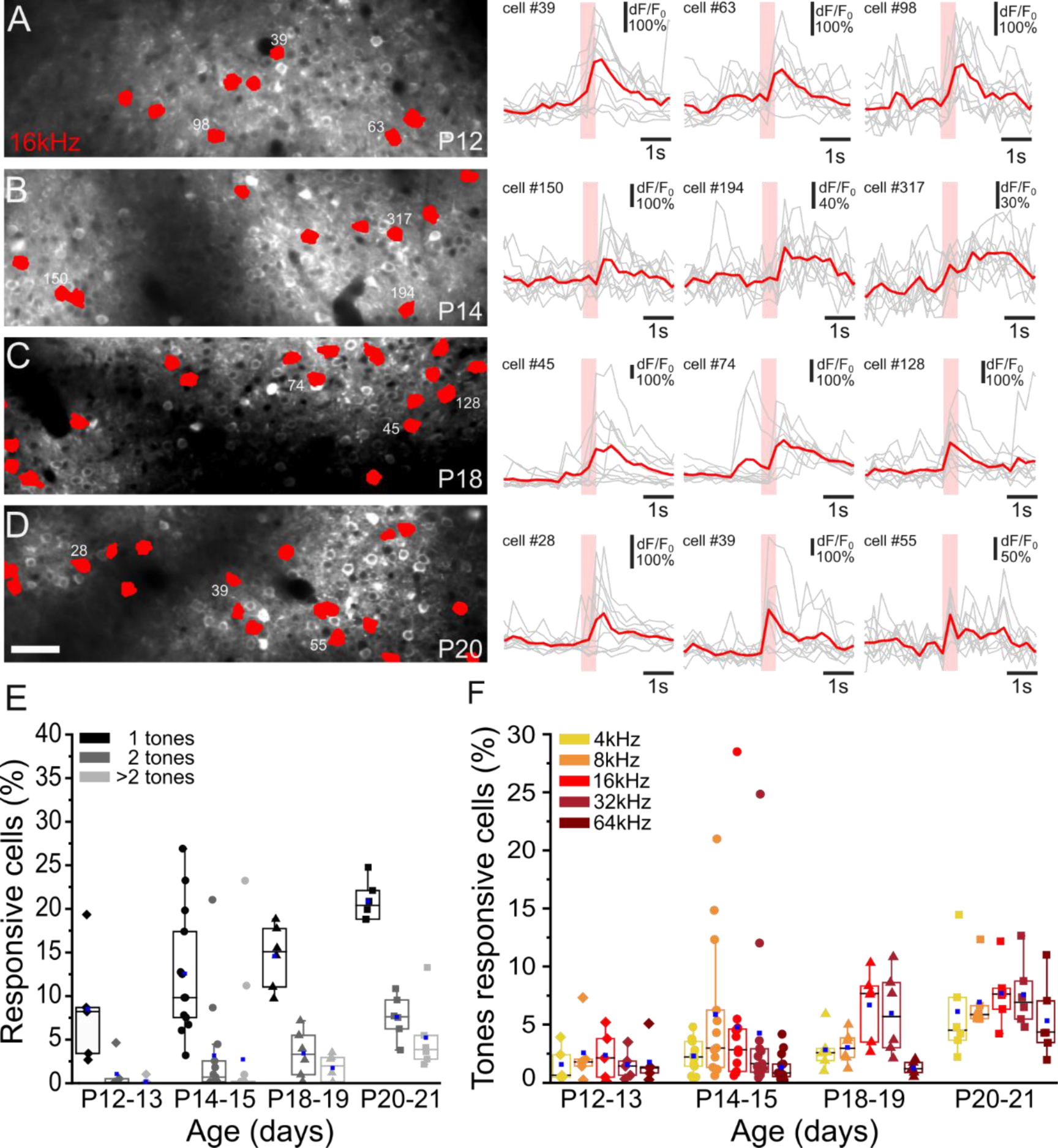
Selective responses to pure tones during development. (A-D) *Left panels*: Examples of fields-of-view showing GCaMP7s expression in ACx L2/3 neurons at P12 (A), P14 (B), P18 (C) and P20 (D). Cells responding to 16kHz tonal stimulus are color-coded in red. Scale bar = 50 µm. *Right panels*: Fluorescence signals (dF/F0) over time from three representative cells responding to 16kHz tonal presentation at different developmental ages. Gray lines represent single trial response. Red lines represent the averaged response to ten stimulus presentation (red rectangle). (E-F) Percentage of cells responding to one tone, two tones or to more then two tones (E) and fraction of cells selectively responding to 4 kHz, 8KHz, 16kHz, 32 kHz and 64kHz (F), in the four developmental windows. For each age the box plots show the median (horizontal black line) and the average value (blue square). No preference toward a tone was identified at P12-P13 (Kruskal-Wallis ANOVA, p = 0.96504, N= 5). P14-P15 and P18-P19 show preferential responses to 4-16kHz or 16-32 kHz respectively (Kruskal-Wallis ANOVA, P12-P13: p = 0.02941, N=13; P18-P19: p = 0.00193, N=6). At P20-P21 no preference was detected (Kruskal-Wallis ANOVA, p = 0.28784).

P14 is a critical age for tonotopic map formation. Thus, we concentrated on this temporal window to better characterize cells’ responses to HF stimuli. At P14-P15 we detected approximately 6% of cells responding to syllables (median, Fig. 2E) whose high frequencies reside between 60 and 90 kHz (Supplementary Fig. 1), whereas only 0.8% of cells responded to 64-kHz pure tone stimulus (median, Fig. 3F). To better characterize P14-P15 responses to HF presentation, we recorded stimulus-evoked activity in response to a set of HF stimuli that resembled syllable characteristics such as frequency range and modulation in time. We exposed P14-P15 mice to four different HF pure tones, namely, 64, 70, 80, and 90 kHz, as well as to increasing and decreasing HF modulated sweeps (70-90 kHz, up-sweep; 90-70 kHz, down-sweep). The analysis of cell coactivation evoked by HF pure tones, up- and down-sweeps, and syllables showed low overlap between different stimuli (Fig. 4A,B). Moreover, the proportion of cells responding to different HF pure tones, up- and down-sweeps, and playback of HF syllables was constant (Kruskal-Wallis ANOVA p= 0,65171, N = 6; Fig. 4A,C), indicating that at P14-P15 a subpopulation of ACx L2/3 cells selectively and specifically responded to more complex features such as frequency modulation, which characterize ultrasound vocal communication.

**Fig. 4.**
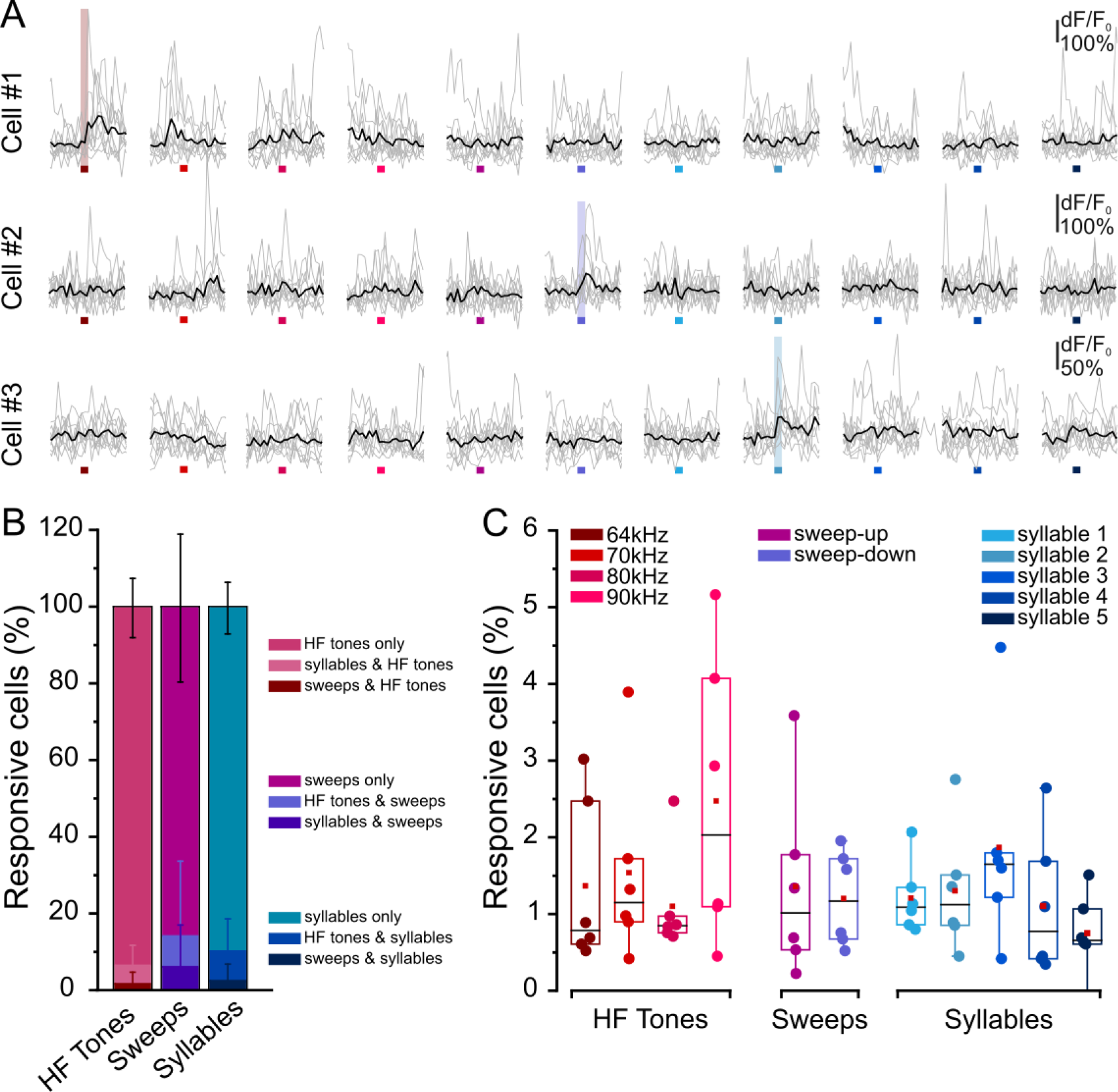
Characterization of ACx L2/3 cells’ responses to high frequency sounds presentation at P14-P15. (A) Fluorescence signals (dF/F_0_) over time from three representative cells selectively responding to 64kHz (cell #1), sweep-down (cell #2) and syllable 2 (cell #3). Stimulus presentation lasts 500 ms. Gray lines represent single trial response to different high-frequency (HF) stimuli (each color represents a stimulus: 64 kHz, 70 kHz, 80 kHz, 90 kHz, sweep-up, sweep-down, syllable 1, syllable 2, syllbale 3, syllable 4, syllable 5). Black lines represent the averaged response to ten stimulus presentation. (B) Fraction of cells selectively responding to “HF tones”, “sweeps” and “sylables” and relative overlapping responses. Values are expressed as average ± sem; N = 6. (C) Percentage of cells selectively responding to: 64 kHz, 70 kHz, 80 kHz, 90 kHz, sweep-up, sweep-down, syllable 1, syllable 2, syllbale 3, syllable 4, syllable 5. Box plots show the median (horizontal black line) and the average value (red square). No difference in the proportion of cells responding to HF pure tones, up- and down-sweeps, and playback of HF syllables was identified (Kruskal-Wallis ANOVA p= 0,65171, N = 6).

### L2/3 cells tuned for the same stimulus displayed highly correlated spontaneous activity

Before hearing onset, developing neural circuits spontaneously generate highly correlated activity in distinct spatial and temporal patterns (Babola et al., 2018; Meng et al., 2021; Wang and Bergles, 2015). These highly stereotyped bursts of action potentials are fundamental for the correct establishment of adult neural circuits that will be later refined through activity-dependent processes upon ear canal opening muller (Clause et al., 2014; Kersbergen et al., 2022; Müller et al., 2019). At hearing onset, sensory-evoked network activation induces increased desynchronization of spontaneous activity and concomitant intensification of sensory-evoked correlation (Meng et al., 2020). To assess how spontaneous (i.e. stimulus-independent) synchronized activity changes during development, we calculated the pairwise correlation between cell pairs in L2/3 of the auditory cortex at P12-P13, P14-P15; P18-P19 and P20-P21, and P20–P21 and evaluated the distribution of correlation coefficients in each developmental window (bin: 0.05; Fig. 5A). Moreover, because pairwise correlated activity can depend on the distance between neurons (Ko et al., 2013; Levy and Reyes, 2012; Liu et al., 2019; Meng et al., 2020; Winkowski and Kanold, 2013), we plotted the average correlations between L2/3 cells as a function of their distance for various developmental windows (bin: 10 µm; Fig. 5B). We found that, as expected, the average pairwise correlation decreased as development proceeded in accordance with the occurrence of desynchronized network activity (Fig. 5A and Supplementary Fig. 2). This increased desynchronization was much more evident for cells located within 150 µm, consistent with previously reported data (Levy and Reyes, 2012; Meng et al., 2020; Winkowski and Kanold, 2013).

Finally, we asked whether cells responding to the same auditory stimulus also displayed a higher spontaneous activity correlation (Fig. 5C-F). Indeed, it was previously demonstrated that, before hearing onset, groups of neurons aligned to the future tonotopic axis that will process similar frequencies of sounds are highly synchronized through the auditory pathway (Babola et al., 2018). To evaluate the relationship between the spontaneous and sound-evoked activity of cells, we looked at the average correlation of spontaneous activity between cell pairs responding to the same stimulus, either pure tones or syllables, and compared it with that obtained from the total cell pairs (including those responding to the same stimulus). At all ages analyzed, the average spontaneous correlation coefficient from cells responding to the same pure tone or syllable resulted to be higher compared to the control group (P12-P13: p = 0,0050084 Wilcoxon rank sum test, “all cells” *vs* “pure tones”; p = 0,0087954, Wilcoxon rank sum test, all cells” *vs* “syllables”; N = 4694, N = 451, N = 326, for “all cells”, “pure tones” and “syllables” respectively; P14-P15: p = 4.891e-89, Wilcoxon rank sum test, “all cells” *vs* “pure tones”; p = 4.8114e-71, Wilcoxon rank sum test, “all cells” *vs* “syllables”; N = 10938, N = 2149, N = 1390, for “all cells”, “pure tones” and “syllables” respectively). Interestingly, this difference was maintained at later developmental windows (P18-P19: p = 1.3129e-44, Wilcoxon rank sum test, “all cells” *vs* “pure tones”; p = 1.6082e-28, Wilcoxon rank sum test, “all cells” *vs* “syllables”; N = 7652, N = 1398, N = 1520, for “all cells”, “pure tones” and “syllables” respectively. P20-P21: p = 4.8953e-20, Wilcoxon rank sum test, “all cells” *vs* “pure tones”; p = 5.0015e-33, Wilcoxon rank sum test, “all cells” *vs* “syllables” N = 3370, N = 1107, N = 958, for “all cells”, “pure tones” and “syllables” respectively), suggesting that groups of neurons processing similar frequencies of sound, that have been shown to exhibit robust correlated activity prior to hearing onset (Babola et al., 2018), maintain this feature also after the opening of the auditory canal when sensory-evoked network activation initiates.

**Fig. 5.**
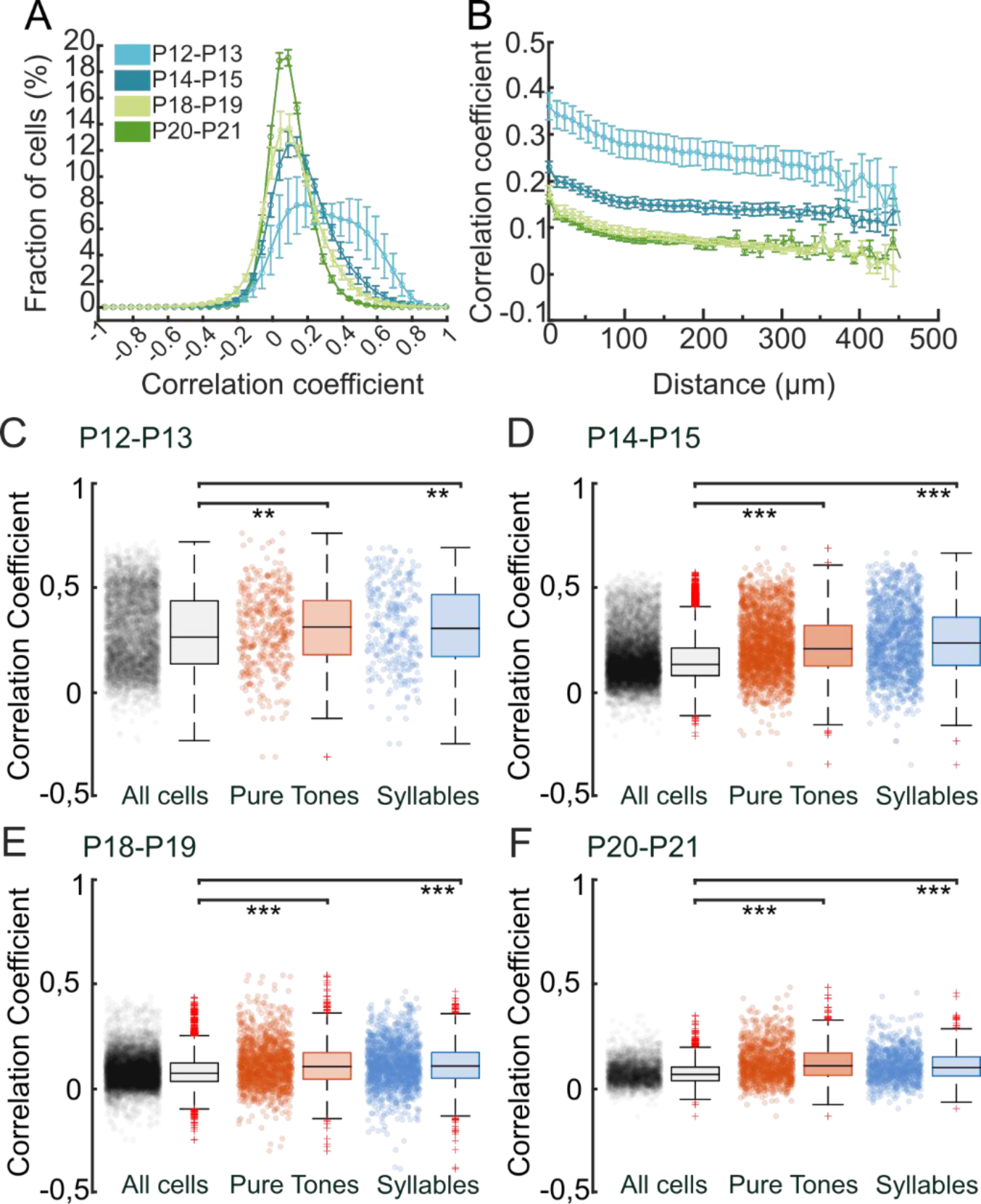
Sound-evoked responses aligh to correlated spontaneous activity. (A-B) Distribution of L2/3 neurons pairwise correlation in ACx (A, bin: 0.05), and average pairwise correlations as a function of cells distance (B, bin: 10 µm) at P12-P13, P14-P15; P18-P19, P20-P21. Cells correlation decreases with age and depends on their distance. Values are expressed as average ± sem. P12-P13 N = 5; P13-P15 N = 13; P18-P19 N = 6; P20-P21 N = 6. (C-F) Distribution of pairwise correlation among L2/3 cells (black) and among cells responsive to pure tones (orange) and syllables (blue) at different developmental ages. Box plots show the median (horizontal black line). Red crosses represent the outliers. **p < 0.01, ***p < 0.001, Wilcoxon rank sum test. P12-P13: N = 4694 (all cells), N = 451 (pure tones), N = 326 (syllables). P14-P15: N = 10938 (all cells), N = 2149 (pure tones), N = 1390 (syllables). P18-P19: N = 7652 (all cells), N = 1398 (pure tones), N = 1520 (syllables). P20-P21: N = 3370 (all cells), N = 1107 (pure tones), N = 958 (syllables).

## Discussion

Many vertebrate species emit sound sequences whose structural complexity communicates information beneficial to the receiver (Gaucher et al., 2013; Lenschow et al., 2022; So et al., 2020; Wang et al., 1995). Mice are no exception and produce high-frequency ultrasound vocalizations relevant to social, sexual, and emotional interactions with conspecifics (Holy and Guo, 2005; Musolf and Penn, 2012). Mouse USVs are auditory signals in the high-frequency range (spanning from 35 to 110 kHz) composed of units of sound called syllables, which differ in their frequency range, structure, and duration (Fonseca et al., 2021). Syllables are arranged in non-random sequences often organized in phrases that have been shown to be predictable of specific behaviors (Agarwalla et al., 2020; Agarwalla et al., 2023; Grimsley et al., 2011). USVs are emitted by pups to elicit maternal retrieval (Ehret and Haack, 1984; Tasaka et al., 2018) and by adults to facilitate social interaction, for example, during courtship behaviors and mate choice (Agarwalla et al., 2023; Lenschow et al., 2022; Musolf et al., 2010; Musolf et al., 2015; White et al., 1998; Yang et al., 2013). Thus, perceiving and encoding USVs plays a prominent role throughout development, from early life up to adulthood. Interestingly, postnatal exposure to acoustic stimuli, including USVs, can impact behavioral choices in adult subjects. For example, adult female mice show a preference for USVs from non-familiar males, but only if they were reared with their father (Asaba et al., 2014; Hammerschmidt et al., 2009; Musolf et al., 2010). Non-familiar USV preference thus requires a learning phase at an early developmental age, during which exposure to father USVs can impact adult auditory processing, similar to what has been described with pure tones and complex sounds exposure, during specific critical developmental windows (Barkat et al., 2011; Insanally et al., 2010; Insanally et al., 2009; Kim and Bao, 2013; Meng et al., 2020; Zhang et al., 2001). Despite the integral role of USV communication in mouse behavior, very little is known about the development of USV hearing capability, and no data are available on when and how USV representation forms during development.

In the present study, we analyzed syllable-evoked activation of layer 2/3 neurons in the auditory cortex of mice in four developmental windows, spanning from hearing onset at the opening of the ear canal (P12-P13), up to P20-P21. Responses to syllables were compared with responses to pure tonal stimuli presentation (4 – 90 kHz) and to HF modulated up- and down-sweeps. Furthermore, by comparing spontaneous activity with sound-evoked responses, we assessed whether L2/3 cell recruitment to the same stimulus at different developmental stages was influenced by previous correlated spontaneous network oscillations.

We observed that syllable-evoked activity in ACx could be detected at hearing onset (P12-P13). The fraction of responding cells to syllables increased during development with a sharp increment at P18-P21. At this age, we also detected a higher fraction of syllable-responding neurons recruited by two or more sound stimuli (either 2 syllables or syllable and pure tone), which were rarely seen at earlier stages but have been previously reported in adult ACx (Carruthers et al., 2013; Kim and Bao, 2013). Thus, compared with pure tone responses, which show a constant increase in the fraction of responsive cells from P12 to P21, and that were described to attain adult-like structure already at P14-P16 in mice (Carrasco et al., 2013; Chen et al., 2021; Polley et al., 2013), USV representation during postnatal development might be slightly delayed, reaching adult-like representation later compared with simple tonal stimuli, consistent with data on the development of complex sounds responses, which have been shown to emerge within a month-long window (Carrasco et al., 2013; Insanally et al., 2010; Insanally et al., 2009; Kim and Bao, 2013).

Previous studies have reported that in both adult rats and mice, neurons in the ACx responding to pure tonal frequencies within the range of USVs are preferentially recruited by the playback of conspecific vocalizations (Calhoun et al., 2023; Carruthers et al., 2013; Kim and Bao, 2013). We found that from P12 to P21, the fraction of cells responding to both syllables and tones increased. This analysis, however, was restricted to tonal stimuli largely below the frequency range that characterizes USVs. Thus, we wondered whether syllable-responding cells could be preferentially activated by pure tones in the frequency range of USVs and/or by up- and down-HF modulated sweeps. At P14-P15, a critical age for tonotopic map formation, we detected low overlap between responses, with most neurons selectively responding to a single category of stimuli (i.e. HF pure tones or HF sweeps or syllables). In accordance with the delayed maturation of complex sound responses and previous studies showing rapid expansion of best frequency responses from lower to higher frequencies between hearing onset and young adult age (Carrasco et al., 2013; Insanally et al., 2010; Insanally et al., 2009; Kim and Bao, 2013; Polley et al., 2013), responses to multiple categories of HF sounds may be delayed and not yet achieved at P14-P15. Furthermore, at P12-P13 and P14-P15, responses to sound presentation were highly variable among animals, coinciding with a dynamic phase for the development of auditory circuits, during which the rapid maturation of cortical responses is strongly susceptible to external manipulation (Barkat et al., 2011; de Villers-Sidani et al., 2007; Insanally et al., 2010; Kim and Bao, 2009; Polley et al., 2013; Zhang et al., 2001; Zhang et al., 2002). The variability across animals was much reduced at P18-P19 and P20-P21, consistent with advanced circuit maturation and achievement of adult-like anatomical organization of ACx subfields (Calhoun et al., 2023; Chen et al., 2021).

The organization of the adult mouse ACx consists of two primary regions surrounded by higher-order areas, mainly responding to USV presentation (Liu et al., 2019; Romero et al., 2020; Stiebler et al., 1997; Tsukano et al., 2015). Interestingly, recent findings in mice identified the left secondary auditory area to exhibit lateralized functional activation and be primarily involved in HF tones and vocalization responses (Calhoun et al., 2023; Geissler and Ehret, 2004; Levy et al., 2019), similar to what has been seen in birds (Schneider and Woolley, 2013) and humans (Norman-Haignere et al., 2015) in processing conspecific communication sounds. Accordingly, for our experiments, we focused on the left ACx; however, how lateralization of USV responses develops and to what extent it is influenced by genetic mechanisms or early experience remains an intriguing future goal.

In the context of cortical development and sensory circuit refinement, spontaneous activity has long been recognized to play an important role in shaping the maturation of sensory systems. This highly synchronized stimulus-independent activity controls numerous aspects of sensory system development, from cell integration to circuit refinement (Kirkby et al., 2013; Leighton and Lohmann, 2016; Luhmann and Khazipov, 2018; Luhmann et al., 2016; Molnár et al., 2020). In the mouse auditory system, spontaneous activity begins prior to hearing onset and continues for almost 2 weeks after birth, when the arrival of auditory signals from the sensory periphery allows neurons to engage in experience-dependent plasticity (Wang and Bergles, 2015). Spontaneous activity originates in the developing cochlea and propagates along the auditory pathway up to the ACx, where it aligns with the future tonotopic axis prior to the initiation of sensory-evoked activity (Babola et al., 2018).

Suppression or manipulation of spontaneous activity significantly impacts the tonotopic precision of auditory neuron connectivity and their tuning properties, as well as the final anatomical organization of brain regions devoted to sound processing (Clause et al., 2014; Kersbergen et al., 2022; Müller et al., 2019), suggesting that spontaneous activity itself contains information that guides fundamental properties of auditory development. Here, we assessed how network coordinated activity is preserved after hearing onset and whether the sensory-dependent maturation of tonotopy contributes to maintaining higher correlated spontaneous activity between neurons responding to the same stimulus.

Our results showed that cell correlation decreased with increasing processing of peripheral auditory stimuli, consistent with previous findings in sensory cortices (Leighton and Lohmann, 2016; Martini et al., 2021). From hearing onset, the “adult-like” desynchronized activity is attended by a progressive increase in incoming auditory stimuli, which refines neuronal responsive properties and reinforces connections between networks involved in processing specific sound features. Our data demonstrate that clusters of neurons responding to the same stimulus maintain a higher correlated spontaneous activity throughout development, even once circuits reach adult-like anatomical and functional organization. This indicates that, while the overall cortical network becomes less synchronized, neurons specialized for the same stimulus feature maintain higher levels of synchronization, which might be responsible for the stronger synaptic connectivity observed in later developmental stages or in mature cortical regions (Ko et al., 2013, 2014; Rossi et al., 2020).

Overall, our data demonstrate for the first time that neurons in the mouse ACx can process complex USVs immediately after the opening of the hearing canal and that their representation increases as the brain develops, with neurons responding to the same stimulus organized in fine-scale sub-networks maintaining highly correlated spontaneous activity after circuits attained mature organization.

Whether USV representation is also linked to higher cognitive functions that help newborn mice associate valence and meaning with stimuli remains to be explored.

## Acknowledgements

We thank the staff of the Laboratory of Adult Neurogeneis at NICO for their constant assistance. The authors thank A. Stella for her support in data analysis and interpretation.

This work was supported by: Human Frontier Science Program (# RGP0003/2020) to S.B., Fondazione Compagnia di San Paolo, Bando Trapezio (#2021.2231) to S.B., Fondazione Umberto Veronesi Postdoctoral Fellowship to C.LR and Marie Skłodowska-Curie Actions to S.Z.

## References

Afrashteh, N., Jafari, Z., Sun, J., Kyweriga, M. and Mohajerani, M.H (2022). Functional organization of mouse auditory cortex in response to stimulus complexity and brain state. bioRxiv doi:10.1101/2022.08.11.503675.

Agarwalla, S., Arroyo, N. S., Long, N. E., O’Brien, W. T., Abel, T. and Bandyopadhyay, S. (2020). Male-specific alterations in structure of isolation call sequences of mouse pups with 16p11.2 deletion. Genes Brain Behav 19, e12681.

Agarwalla, S., De, A. and Bandyopadhyay, S. (2023). Predictive Mouse Ultrasonic Vocalization Sequences: Uncovering Behavioral Significance, Auditory Cortex Neuronal Preferences, and Social-Experience-Driven Plasticity. J Neurosci 43, 6141–6163.

Anthwal, N. and Thompson, H. (2016). The development of the mammalian outer and middle ear. J Anat 228, 217–232.

Arriaga, G. and Jarvis, E. D. (2013). Mouse vocal communication system: are ultrasounds learned or innate? Brain Lang 124, 96–116.

Asaba, A., Okabe, S., Nagasawa, M., Kato, M., Koshida, N., Osakada, T., Mogi, K. and Kikusui, T. (2014). Developmental social environment imprints female preference for male song in mice. PLoS One 9, e87186.

Babola, T. A., Li, S., Gribizis, A., Lee, B. J., Issa, J. B., Wang, H. C., Crair, M. C. and Bergles, D. E. (2018). Homeostatic Control of Spontaneous Activity in the Developing Auditory System. Neuron 99, 511–524.e515.

Barkat, T. R., Polley, D. B. and Hensch, T. K. (2011). A critical period for auditory thalamocortical connectivity. Nat Neurosci 14, 1189–1194.

Calhoun, G., Chen, C. T. and Kanold, P. O. (2023). Bilateral widefield calcium imaging reveals circuit asymmetries and lateralized functional activation of the mouse auditory cortex. Proc Natl Acad Sci U S A 120, e2219340120.

Carrasco, M. M., Trujillo, M. and Razak, K. (2013). Development of response selectivity in the mouse auditory cortex. Hear Res 296, 107–120.

Carruthers, I. M., Natan, R. G. and Geffen, M. N. (2013). Encoding of ultrasonic vocalizations in the auditory cortex. J Neurophysiol 109, 1912–1927.

Chabout, J., Sarkar, A., Dunson, D. B. and Jarvis, E. D. (2015). Male mice song syntax depends on social contexts and influences female preferences. Front Behav Neurosci 9, 76.

Chen, F., Takemoto, M., Nishimura, M., Tomioka, R. and Song, W. J. (2021). Postnatal development of subfields in the core region of the mouse auditory cortex. Hear Res 400, 108138.

Chen, T. W., Wardill, T. J., Sun, Y., Pulver, S. R., Renninger, S. L., Baohan, A., Schreiter, E. R., Kerr, R. A., Orger, M. B., Jayaraman, V., et al. (2013). Ultrasensitive fluorescent proteins for imaging neuronal activity. Nature 499, 295–300.

Clause, A., Kim, G., Sonntag, M., Weisz, C. J., Vetter, D. E., Rűbsamen, R. and Kandler, K. (2014). The precise temporal pattern of prehearing spontaneous activity is necessary for tonotopic map refinement. Neuron 82, 822–835.

Dana, H., Sun, Y., Mohar, B., Hulse, B. K., Kerlin, A. M., Hasseman, J. P., Tsegaye, G., Tsang, A., Wong, A., Patel, R., et al. (2019). High-performance calcium sensors for imaging activity in neuronal populations and microcompartments. Nat Methods 16, 649–657.

de Villers-Sidani, E., Chang, E. F., Bao, S. and Merzenich, M. M. (2007). Critical period window for spectral tuning defined in the primary auditory cortex (A1) in the rat. J Neurosci 27, 180–189.

Ehret, G. (1983). Development of hearing and response behavior to sound stimuli: behavioral studies. R. Romand (Ed.), Development of auditory and vestibular systems, Academic Press, New York (1983), pp. 211–237.

Ehret, G. and Haack, B. (1984). Motivation and arousal influence sound-induced maternal pup-retrieving behavior in lactating house mice. Z Tierpsychol 65:25–39.

Fonseca, A. H., Santana, G. M., Bosque Ortiz, G. M., Bampi, S. and Dietrich, M. O. (2021). Analysis of ultrasonic vocalizations from mice using computer vision and machine learning. Elife 10.

Garcia-Lazaro, J. A., Shepard, K. N., Miranda, J. A., Liu, R. C. and Lesica, N. A. (2015). An Overrepresentation of High Frequencies in the Mouse Inferior Colliculus Supports the Processing of Ultrasonic Vocalizations. PLoS One 10, e0133251.

Gaucher, Q., Huetz, C., Gourévitch, B., Laudanski, J., Occelli, F. and Edeline, J. M. (2013). How do auditory cortex neurons represent communication sounds? Hear Res 305, 102–112.

Geissler, D. B. and Ehret, G. (2004). Auditory perception vs. recognition: representation of complex communication sounds in the mouse auditory cortical fields. Eur J Neurosci 19, 1027–1040.

Grimsley, J. M., Monaghan, J. J. and Wenstrup, J. J. (2011). Development of social vocalizations in mice. PLoS One 6, e17460.

Hammerschmidt, K., Radyushkin, K., Ehrenreich, H. and Fischer, J. (2009). Female mice respond to male ultrasonic ’songs’ with approach behaviour. Biol Lett 5, 589–592.

Holy, T. E. and Guo, Z. (2005). Ultrasonic songs of male mice. PLoS Biol 3, e386.

Insanally, M. N., Albanna, B. F. and Bao, S. (2010). Pulsed noise experience disrupts complex sound representations. J Neurophysiol 103, 2611–2617.

Insanally, M. N., Köver, H., Kim, H. and Bao, S. (2009). Feature-dependent sensitive periods in the development of complex sound representation. J Neurosci 29, 5456–5462.

Jouhaneau, J. and Bagady, A. (1984). Effect of early auditory stimulation on the choice of acoustical environment by adult Swiss albino mice (Mus musculus). J Comp Psychol 98, 318–326.

Kersbergen, C. J., Babola, T. A., Rock, J. and Bergles, D. E. (2022). Developmental spontaneous activity promotes formation of sensory domains, frequency tuning and proper gain in central auditory circuits. Cell Rep 41, 111649.

Kim, H. and Bao, S. (2009). Selective increase in representations of sounds repeated at an ethological rate. J Neurosci 29, 5163–5169.

Kim, H. and Bao, S. (2013). Experience-dependent overrepresentation of ultrasonic vocalization frequencies in the rat primary auditory cortex. J Neurophysiol 110, 1087–1096.

Kirkby, L. A., Sack, G. S., Firl, A. and Feller, M. B. (2013). A role for correlated spontaneous activity in the assembly of neural circuits. Neuron 80, 1129–1144.

Ko, H., Cossell, L., Baragli, C., Antolik, J., Clopath, C., Hofer, S. B. and Mrsic-Flogel, T. D. (2013). The emergence of functional microcircuits in visual cortex. Nature 496, 96–100.

Ko, H. (2014). Functional organization of synaptic connections in the neocortex. Science. 31;346(6209):555.

Kronman, F. A., Liwang, J. K., Betty, R., Vanselow, D. J., Wu, Y., Tustison, N. J., Bhandiwad, A., Manjila, S. B., Minteer, J. A., Shin, D., Lee, C. H., Patil, R., Duda, J. T., Puelles, L., Gee, J. C., Zhang, J., Ng, L. and Kim, Y. (2023). Developmental Mouse Brain Common Coordinate Framework. bioRxiv doi: 10.1101/2023.09.14.557789.

Leighton, A. H. and Lohmann, C. (2016). The Wiring of Developing Sensory Circuits-From Patterned Spontaneous Activity to Synaptic Plasticity Mechanisms. Front Neural Circuits 10, 71.

Lenschow, C., Mendes, A. R. P. and Lima, S. Q. (2022). Hearing, touching, and multisensory integration during mate choice. Front Neural Circuits 16, 943888.

Levy, R. B., Marquarding, T., Reid, A. P., Pun, C. M., Renier, N. and Oviedo, H. V. (2019). Circuit asymmetries underlie functional lateralization in the mouse auditory cortex. Nat Commun 10, 2783.

Levy, R. B. and Reyes, A. D. (2012). Spatial profile of excitatory and inhibitory synaptic connectivity in mouse primary auditory cortex. J Neurosci 32, 5609–5619.

Liu, J., Whiteway, M. R., Sheikhattar, A., Butts, D. A., Babadi, B. and Kanold, P. O. (2019). Parallel Processing of Sound Dynamics across Mouse Auditory Cortex via Spatially Patterned Thalamic Inputs and Distinct Areal Intracortical Circuits. Cell Rep 27, 872–885.e877.

Luhmann, H. J. and Khazipov, R. (2018). Neuronal activity patterns in the developing barrel cortex. Neuroscience 368, 256–267.

Luhmann, H. J., Sinning, A., Yang, J. W., Reyes-Puerta, V., Stüttgen, M. C., Kirischuk, S. and Kilb, W. (2016). Spontaneous Neuronal Activity in Developing Neocortical Networks: From Single Cells to Large-Scale Interactions. Front Neural Circuits 10, 40.

Martini F. J., Guillamón-Vivancos T., Moreno-Juan V., Valdeolmillos M. and López-Bendito G. (2021) Spontaneous activity in developing thalamic and cortical sensory networks. Neuron 109, 2519–2534.

McMullen, N. T., Goldberger, B. and Glaser, E. M. (1988). Postnatal development of lamina III/IV nonpyramidal neurons in rabbit auditory cortex: quantitative and spatial analyses of Golgi-impregnated material. J Comp Neurol 278, 139–155.

Meng, X., Mukherjee, D., Kao, J. P. Y. and Kanold, P. O. (2021). Early peripheral activity alters nascent subplate circuits in the auditory cortex. Sci Adv 7.

Meng, X., Solarana, K., Bowen, Z., Liu, J., Nagode, D. A., Sheikh, A., Winkowski, D. E., Kao, J. P. Y. and Kanold, P. O. (2020). Transient Subgranular Hyperconnectivity to L2/3 and Enhanced Pairwise Correlations During the Critical Period in the Mouse Auditory Cortex. Cereb Cortex 30, 1914–1930.

Mikaelian D, Ruben RJ (1965). Development of hearing in the normal Cba-J mouse: correlation of physiological observations with behavioral responses and with cochlear anatomy. Acta Otolaryngol 59: 451–461.

Molnár, Z., Luhmann, H. J. and Kanold, P. O. (2020). Transient cortical circuits match spontaneous and sensory-driven activity during development. Science 370.

Musolf, K., Hoffmann, F. and Penn, D. J. (2010). Ultrasonic courtship vocalizations in wild house mice, Mus musculus musculus pp. 757–764. Animal Behaviour: Elsevier.

Musolf, K. and Penn, D. J. (2012). Ultrasonic vocalizations in house mice: a cryptic mode of acoustic communication. In: Macholan, Milos; Baird, Stuart J E; Munclinger, Pavel; Pialek, Jaroslaw. Evolution of the house mouse. Cambridge: Cambridge University Press, 253–277.

Musolf, K., Meindl, S., Larsen, A. L., Kalcounis-Rueppell, M. C. and Penn, D. J. (2015). Ultrasonic Vocalizations of Male Mice Differ among Species and Females Show Assortative Preferences for Male Calls. PLoS One 10, e0134123.

Müller, N. I. C., Sonntag, M., Maraslioglu, A., Hirtz, J. J. and Friauf, E. (2019). Topographic map refinement and synaptic strengthening of a sound localization circuit require spontaneous peripheral activity. J Physiol 597, 5469–5493.

Norman-Haignere, S., Kanwisher, N. G. and McDermott, J. H. (2015). Distinct Cortical Pathways for Music and Speech Revealed by Hypothesis-Free Voxel Decomposition. Neuron 88, 1281–1296.

Pachitariu, M., Stringer, C., Dipoppa, M., Schröder, S., Rossi, L. F., Dalgleish, H., Carandini, M. and Harris K. D. (2017). Suite2p: beyond 10,000 neurons with standard two-photon microscopy. bioRxiv doi: 10.1101/061507.

Polley, D. B., Thompson, J. H. and Guo, W. (2013). Brief hearing loss disrupts binaural integration during two early critical periods of auditory cortex development. Nat Commun 4, 2547.

Romero, S., Hight, A. E., Clayton, K. K., Resnik, J., Williamson, R. S., Hancock, K. E. and Polley, D. B. (2020). Cellular and Widefield Imaging of Sound Frequency Organization in Primary and Higher Order Fields of the Mouse Auditory Cortex. Cereb Cortex 30, 1603–1622.

Rossi, L.F., Harris, K.D. and Carandini, M. (2020). Spatial connectivity matches direction selectivity in visual cortex. Nature. 588, 648–652.

Sangiamo, D. T., Warren, M. R. and Neunuebel, J. P. (2020). Ultrasonic signals associated with different types of social behavior of mice. Nat Neurosci 23, 411–422.

Scattoni, M. L., Ricceri, L. and Crawley, J. N. (2011). Unusual repertoire of vocalizations in adult BTBR T+tf/J mice during three types of social encounters. Genes Brain Behav 10, 44–56.

Schachtele, S. J., Losh, J., Dailey, M. E. and Green, S. H. (2011). Spine formation and maturation in the developing rat auditory cortex. J Comp Neurol 519, 3327–3345.

Schiavo, J. K., Valtcheva, S., Bair-Marshall, C. J., Song, S. C., Martin, K. A. and Froemke, R. C. (2020). Innate and plastic mechanisms for maternal behaviour in auditory cortex. Nature 587, 426–431.

Schneider, D. M. and Woolley, S. M. (2013). Sparse and background-invariant coding of vocalizations in auditory scenes. Neuron 79, 141–152.

So, N. L. T., Edwards, J. A. and Woolley, S. M. N. (2020). Auditory Selectivity for Spectral Contrast in Cortical Neurons and Behavior. J Neurosci 40, 1015–1027.

Stiebler, I., Neulist, R., Fichtel, I. and Ehret, G. (1997). The auditory cortex of the house mouse: left-right differences, tonotopic organization and quantitative analysis of frequency representation. J Comp Physiol A 181, 559–571.

Tasaka, G. I., Guenthner, C. J., Shalev, A., Gilday, O., Luo, L. and Mizrahi, A. (2018). Genetic tagging of active neurons in auditory cortex reveals maternal plasticity of coding ultrasonic vocalizations. Nat Commun 9, 871.

Tsukano, H., Horie, M., Bo, T., Uchimura, A., Hishida, R., Kudoh, M., Takahashi, K., Takebayashi, H. and Shibuki, K. (2015). Delineation of a frequency-organized region isolated from the mouse primary auditory cortex. J Neurophysiol 113, 2900–2920.

Wang, H. C. and Bergles, D. E. (2015). Spontaneous activity in the developing auditory system. Cell Tissue Res 361, 65–75.

Wang, X., Merzenich, M. M., Beitel, R. and Schreiner, C. E. (1995). Representation of a species-specific vocalization in the primary auditory cortex of the common marmoset: temporal and spectral characteristics. J Neurophysiol 74, 2685–2706.

White, N. R., Prasad, M., Barfield, R. J. and Nyby, J. G. (1998). 40- and 70-kHz vocalizations of mice (Mus musculus) during copulation. Physiol Behav 63, 467–473.

Winkowski, D. E. and Kanold, P. O. (2013). Laminar transformation of frequency organization in auditory cortex. J Neurosci 33, 1498–1508.

Yang, E. J., Lin, E. W. and Hensch, T. K. (2012). Critical period for acoustic preference in mice. Proc Natl Acad Sci U S A 109 Suppl 2, 17213–17220.

Yang, M., Loureiro, D., Kalikhman, D. and Crawley, J. N. (2013). Male mice emit distinct ultrasonic vocalizations when the female leaves the social interaction arena. Front Behav Neurosci 7, 159.

Zhang, L. I., Bao, S. and Merzenich, M. M. (2001). Persistent and specific influences of early acoustic environments on primary auditory cortex. Nat Neurosci 4, 1123–1130.

Zhang, L. I., Bao, S. and Merzenich, M. M. (2002). Disruption of primary auditory cortex by synchronous auditory inputs during a critical period. Proc Natl Acad Sci U S A 99, 2309–2314.

Zucca, S., D’Urso, G., Pasquale, V., Vecchia, D., Pica, G., Bovetti, S., Moretti, C., Varani, S., Molano-Mazón, M., Chiappalone, M., et al. (2017). An inhibitory gate for state transition in cortex. Elife 6.

