## Supplementary Information for "Developmental encoding of ultrasound vocalizations in the mouse auditory cortex"

**Supplementary Figure 1**

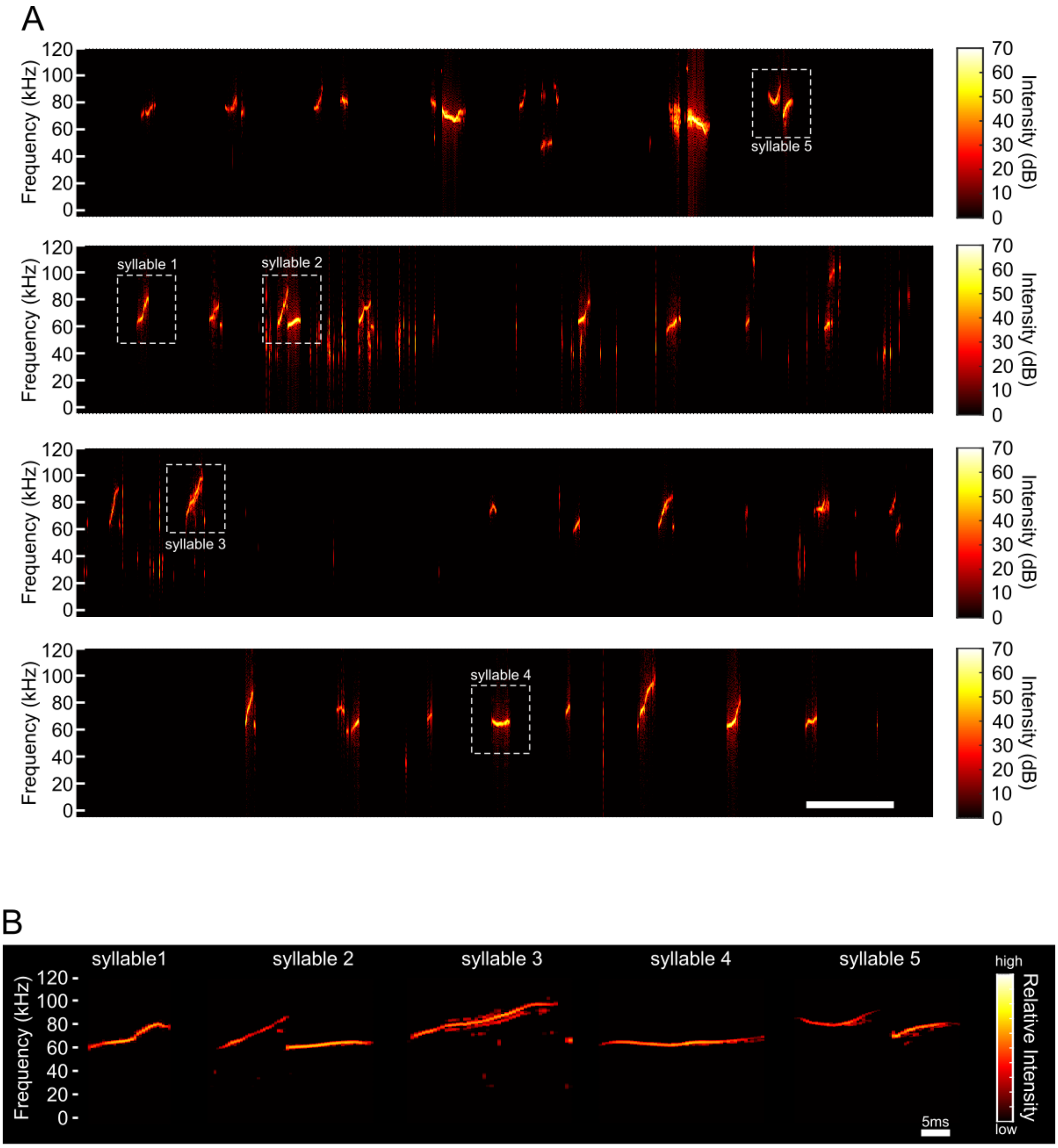


**Supplementary Fig. 1. Ultra-sound vocalizations (USVs) emitted by a C57BL6J male mouse.** (A) Four examples of USVs emitted by an adult (P60) C57BL6J male mouse when exposed to female urine. USVs were recorded at a sampling frequency of 200 kHz via an ultrasound microphone CM16/CMPA (Avisoft Bioacoustic, Germany), coupled with a recording interface (UltrasoundGate 116Hb, Avisoft Bioacoustic) and recording software (Avisoft Recorder). All recordings were performed in a sound-proof box and USVs were elicited by exposing adult male mice to females' scent marks. (B) Five syllables differing in their length, frequency, and shape were isolated from USV recordings.

**Supplementary Figure 2**

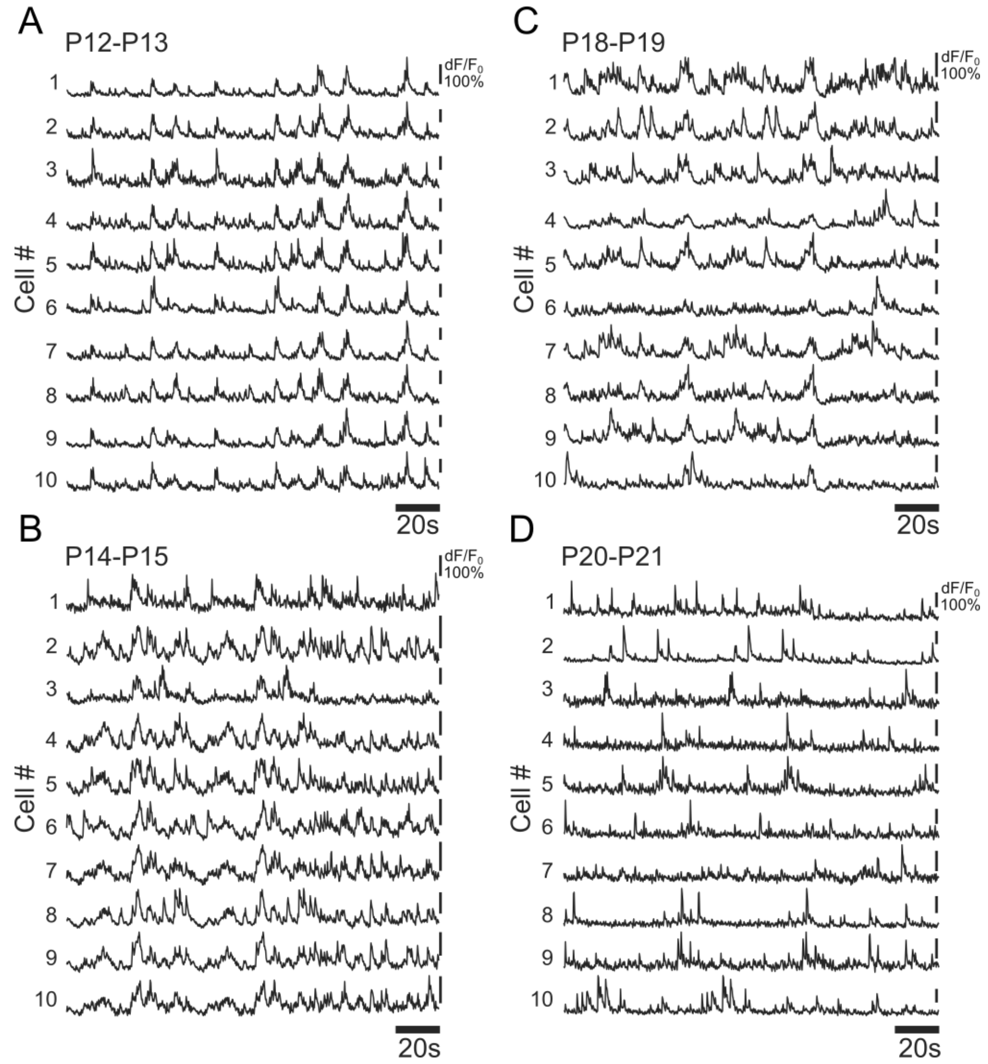


**Supplementary Fig. 2. Spontaneous activity in L2/3 neurons desynchronizes while development proceeds.** (A-D) Spontaneous fluorescence signals (dF/$F_0$) over time from 10 representative L2/3 ACx cells, at P12-P13 (A), P14-P15 (B), P18-P19 (C) an
